# A long-lived pool of PINK1 imparts a molecular memory of depolarisation-induced activity

**DOI:** 10.1101/2024.07.03.601901

**Authors:** Liam Pollock, Ioanna Ch. Georgiou, Emma V. Rusilowicz-Jones, Michael J. Clague, Sylvie Urbé

**Author notes:** These authors contributed equally (Liam Pollock, Ioanna Ch. Georgiou).

## Abstract

The Parkinson’s disease linked kinase, PINK1, is a short lived protein that undergoes cleavage upon mitochondrial import leading to its release to the cytosol and proteasomal degradation. Under mitochondria depolarising conditions, it accumulates on mitochondria where it becomes activated, phosphorylating both ubiquitin and the ubiquitin E3 ligase Parkin, at Ser65. Here we have used a ubiquitylation inhibitor TAK-243 to accumulate cleaved PINK1 (cPINK1) in a cell line that lacks Parkin. We show that cPINK1 phosphorylates free ubiquitin and can be released to the cytosol in an active form. We show that in RPE1 cells under mitochondria depolarising conditions (i) the majority of PINK1 cleavage proceeds unimpeded and (ii) accrued PINK1 cannot be accounted for by protein stabilisation alone. Accordingly, we suggest that translation of PINK1 mRNA must be mobilised under mitochondrial depolarisation. We have further discovered a pre-conditioning phenomenon, whereby an initial depolarising treatment leaves a residual pool of active PINK1, which remains competent for seeding the activation of nascent cPINK1, despite a 16 hour recuperation period.

## Introduction

PINK1 is a Parkinson’s disease-linked kinase which selectively accumulates at impaired or damaged mitochondria [1–3]. Here it phosphorylates Ser 65 on both ubiquitin (hereafter pUb) and the E3 ubiquitin ligase Parkin (PRKN) [4–7]. This activates Parkin, sparking a feed-forward cascade, resulting in widespread ubiquitylation at the mitochondrial surface, leading to mitophagy [8, 9]. The accepted model suggests that PINK1 is constitutively expressed and imported into mitochondria via the outer mitochondrial import (TOM) complex. It then engages with the inner mitochondrial import machinery and undergoes cleavage within its transmembrane domain by the mitochondrial inner membrane rhomboid protease presenilin-associated rhomboid-like protein (PARL) [1, 10, 11]. The cleaved form of PINK1 (cPINK1) is then released from the mitochondria and rapidly degraded by the proteasome [12, 13]. Loss of the mitochondrial membrane potential prompts uncleaved membrane anchored PINK1 to accumulate in association with the TOM complex on the outer mitochondrial membrane, with the kinase domain facing the cytosol [14, 15]. This PINK1 stabilisation requires the small TOM complex subunit TOM7 [16, 17].

In the presence of proteasome inhibitors, the cleaved form of PINK1 accumulates in the cytosol [12]. N-terminal truncated forms of PINK1 mimic this cleavage and show residual kinase activity towards ubiquitin [18]. Moreover, the kinase activity of truncated PINK1 can protect neurons from the neurotoxin MPTP [19] and can inhibit global translation via phosphorylation of eEF1A [20]. The influence of PINK1 kinase activity extends to the nucleus and several proteins involved in nuclear confined processes acquire pSer65 ubiquitylation following mitochondrial depolarisation in a PINK1-dependent manner [21–23]. Whether or not these modifications require a freely diffusible form of PINK1 is currently unclear. Whilst accumulation of full length PINK1 at mitochondria provides a sensor for mitochondrial stress, the accumulation of cytosolic cleaved PINK1 may provide an analogous sensing mechanism for proteasomal stress. Concordantly the truncated form of PINK1 can protect against cell death induced by proteasome inhibition [24, 25].

In neurons, the transit to and residence time within synaptic terminals far exceeds the lifetime of PINK1. To preserve PINK1-dependent quality control at nerve terminals, PINK1 mRNA attaches to the mitochondrial outer membrane proteins synaptojanin 2 binding protein (SYNJ2BP) and synaptojanin 2 (SYNJ2) [26]. Furthermore PINK1 has been linked to the TOM20-dependent mitochondrial localisation and Parkin-mediated translational derepression of discrete nuclear encoded respiratory chain complex proteins [27].

In this paper we have used TAK-243, an inhibitor of the E1 ubiquitin ligase UBA1, to accumulate the cleaved form of PINK1 in an RPE1 cell line that naturally lacks Parkin [28]. We find that cPINK1 continues to be generated at the same basal rate under mitochondrial depolarising conditions, whilst a pathway to full length PINK1 is concurrently activated. Our data suggest this accumulation of full length PINK1 cannot be fully accounted for by stabilisation of PINK1 alone and likely reflects an element of increased translation. TAK-243 application, during recovery from a mitochondria depolarising treatment, allows us to compare the rate at which conjugated pUb is lost alongside total ubiquitin. Notably, we discover a second wave of free pUb generation, coincident with the emergence of cPINK1. On investigating recovery of cells from mitochondrial depolarisation, we find that a prior depolarising treatment elicits phosphorylation of cleaved PINK1 generated in response to TAK-243, 16 hours after removal of the depolarising agents. We propose a model in which PINK1 accumulation, under depolarising conditions, in part reflects mobilisation of its own translation at mitochondria and that a stable residual pool of pre-activated PINK1, conditions mitochondria for the activation of nascent PINK1.

## Results

### TAK-243 redirects PINK1 activity towards free ubiquitin

Throughout this study, we have utilised TAK-243 to inhibit the ubiquitin E1 enzyme UBA1, the principal initiating enzyme of ubiquitylation cascades [28–30]. Its application results in the rapid downshift in molecular weight of UBA1 consistent with a loss of charged ubiquitin in HCT116 and hTERT1-RPE1 cells. A more gradual loss of ubiquitin from the E2 enzyme UBC13 (UBE2N) is also observed (Figure 1A). Acute application of TAK-243 can be used to visualise the dissipation of global ubiquitylation within hours in RPE1 cells (Figure 1B). We compared the pUb Western blotting profile of cells subjected to mitochondrial depolarisation by the application of the respiratory chain inhibitors Antimycin A and Oligomycin A (AO), in the presence and absence of TAK-243. Existing mitochondrial ubiquitin can serve as a PINK1 substrate, but the pUb signal is highly amplified by depolarisation-induced ubiquitylation even though these cells do not express Parkin (Figure 1C). In the presence of TAK-243, the complexity of the pUb profile following mitochondrial depolarisation is reduced. A few higher molecular weight bands become more prominent against this reduced background and are candidates for “priming” substrates that enable the subsequent cascade [31]. Notably when ubiquitylation is inhibited, PINK1 directs its activity towards free ubiquitin molecules (Figure 1C-E). The distribution of pUb within subcellular fractions is shown in Figure 1D and E. pUb conjugation is clearly not restricted to mitochondria and the population of free pUb is highly enriched in the cytosolic fraction. SDS-PAGE is routinely carried out under reducing conditions which liberates E1 and E2 associated ubiquitin as well as any otherwise thioester linked ubiquitin. Under non-reducing conditions we retain free pUb confirming that this is part of the free ubiquitin pool rather than loaded onto conjugating enzymes (Figure 1E).

**Figure 1.**
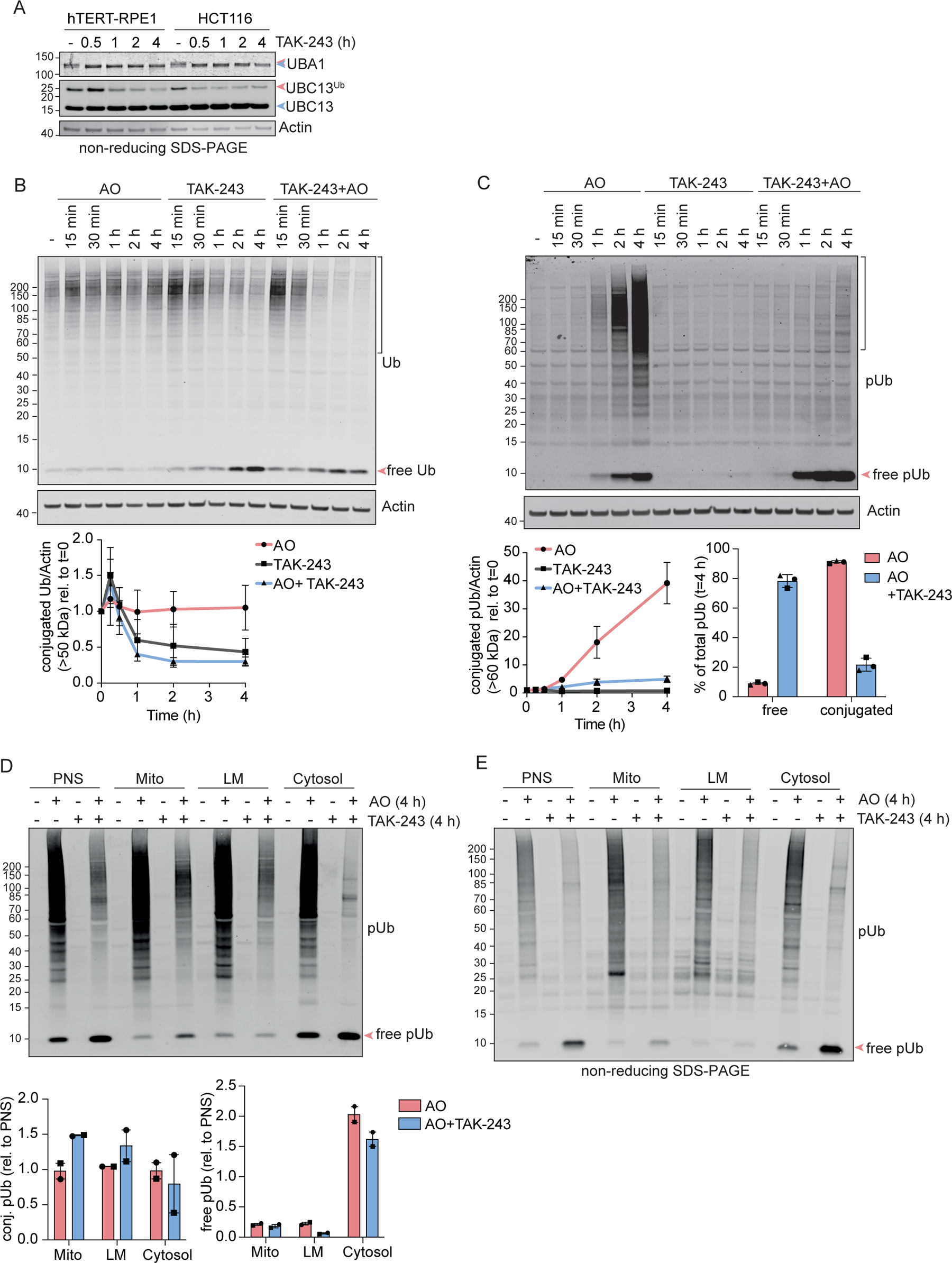
The UBA1 inhibitor TAK-243 reveals de novo phosphorylation of free ubiquitin in the cytosol. **A.** hTERT-RPE1 Flp-In and HCT116 cells were treated ± TAK-243 (1 μM) for the indicated times, lysed and proteins analysed by non-reducing SDS-PAGE and western blot. Blue and red arrowheads indicate the unloaded forms and the Ub-loaded forms respectively of UBA1 and UBC13. **B and C**. hTERT-RPE1 Flp-In cells were treated ± TAK-243 (1 μM), Antimycin A (1 μM) and Oligomycin A (1 μM) (AO), or both TAK-243 and AO. Representative western blot of three independent experiments and quantification showing the decay of conjugated ubiquitin (B) and accumulation of free and conjugated pSer65-Ubiquitin (pUb, C). Brackets indicate the MW range used for the quantification. Error bars show standard deviation. **D and E.** hTERT-RPE1 Flp-In cells were treated for 4 h ± TAK-243 (1 μM), AO (1 μM), or TAK-243 and AO and subjected to subcellular fractionation. PNS: Post-nuclear supernatant; Mito: mitochondria enriched fraction; LM: light membrane fraction. TOMM20 serves as a mitochondrial marker. SDS-PAGE was performed under reducing (D) or non-reducing (E) conditions to assess levels of free pUb. Bracket indicates the MW range of signal used for conjugated pUb quantification. Quantification of data shown in D shows the enrichment of conjugated and free pUb in mitochondrial, light membrane and cytosolic fractions (normalised to PNS) for two independent experiments. Error bars show the range.

### TAK-243 treatment stabilises a cleaved form of PINK1

The application of TAK-243 can be functionally equivalent to proteasome inhibition in terms of protein stabilisation. We compared the effects of treatment with AO, TAK-243, or the proteasome inhibitor Epoxomicin, alone or in combination. Here we confirm that TAK-243 leads to the appearance of a cleaved form of PINK1 (cPINK1), exactly as previously reported with proteasome inhibitors (Figure 2A) [3, 12, 32]. Both Epoxomicin and TAK-243 generate similar amounts of cPINK1 (Figure 2B). Remarkably the acquisition rate of this cPINK1, is completely unaffected by concomitant AO treatment, which instead leads to the parallel acquisition of full length PINK1. The full length PINK1 that accumulates under depolarising conditions, is isolated from this cleavage pathway and accrues at the same rate ± UBA1 inhibition (Figure 2A). As expected, full length PINK1 is confined to mitochondria, whilst a significant fraction of cPINK1 appears in the cytosolic fraction (Figure 2C). We next accumulated full length PINK1, through a long period of depolarisation (24 hours) and added TAK-243 for the final 2 hours. cPINK1 is generated upon TAK-243 application, at the same rate ± mitochondrial depolarisation (Figure 2D). Note that this experiment also reveals a minor fraction of mono-ubiquitylated full length PINK1 which is lost within the 2 hours of TAK-243 treatment. To demonstrate that the TAK-243 induced cPINK1 requires mitochondria we used a method to remove mitochondria from cells in advance of drug application [33, 34]. RPE1 cells stably expressing an excess of YFP-Parkin are first treated with AO for 24 hours which removes mitochondria through mitophagy. In these mitochondria deficient cells, cPINK1 no longer accumulates following TAK-243 application (Figure 2E).

**Figure 2.**
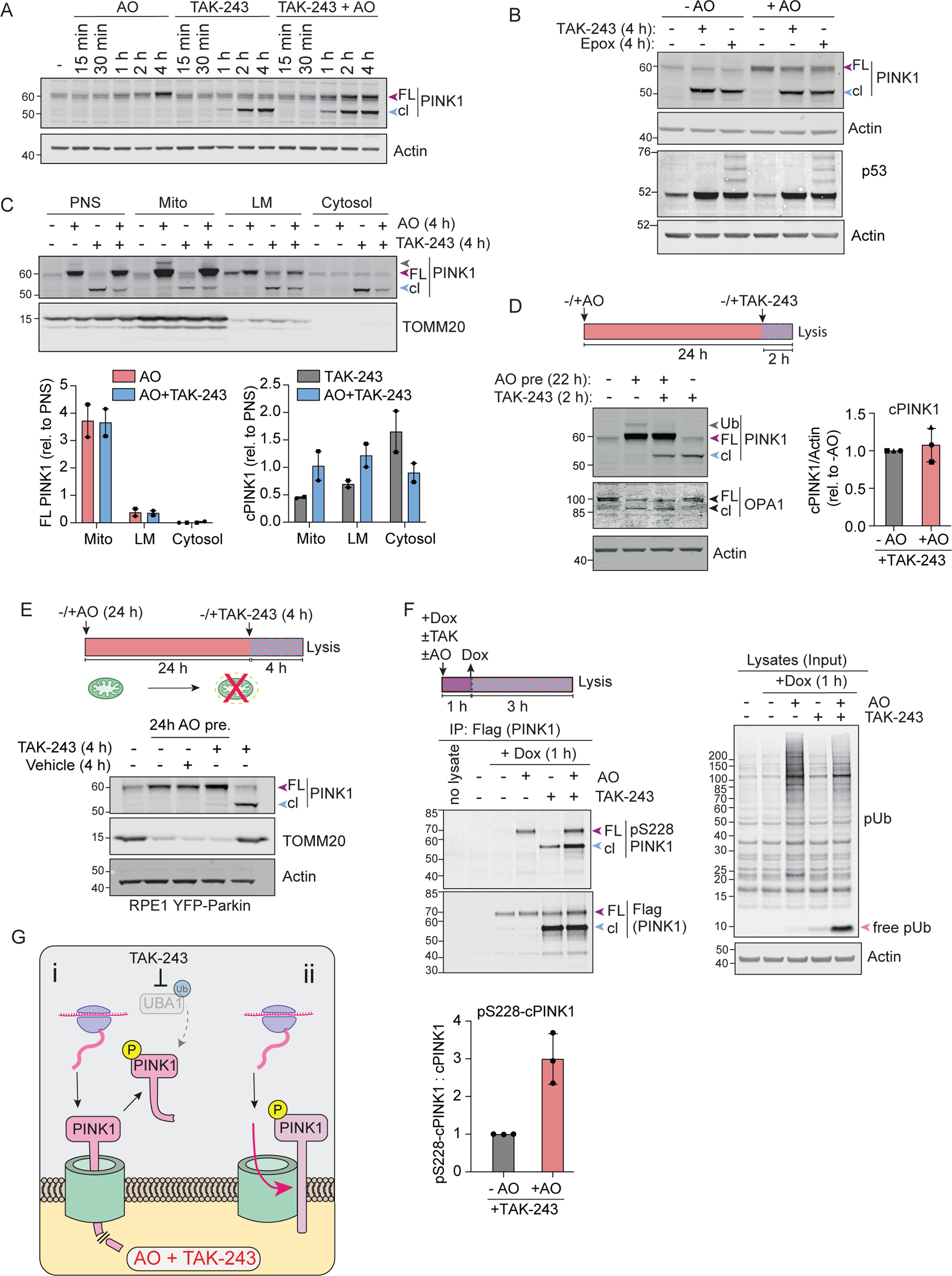
TAK-243 reveals continued processing of PINK1 and generation of cleaved active PINK1 at depolarised mitochondria. **A.** hTERT-RPE1 Flp-In cells were treated ± TAK-243 (1 μM), Antimycin A (1 μM) and Oligomycin A (1 μM) (AO), or both TAK-243 and AO. Representative western blots of three independent experiments. Coloured arrowheads correspond to full length (FL) and cleaved PINK1 (cl). **B.** hTERT-RPE1 Flp-In cells were treated for 4 h ±AO (1 μM each), ± Epoxomicin (100 nM) or ± TAK-243 (1 μM). Representative western blot of two independent experiments is shown. **C**. hTERT-RPE1 Flp-In cells were treated ± AO, or TAK-243 and AO (1 μM each) for 4 h before subcellular fractionation (same samples as in Figure 1D). Representative western blot of two independent experiments and associated quantification of the relative enrichment of full length (FL) and cleaved PINK1 (cPINK1) are shown. PNS: Post-nuclear supernatant; Mito: mitochondria enriched fraction; LM: light membrane fraction. TOMM20 serves as a mitochondrial marker. Error bars show the range. **D.** hTERT-RPE1 Flp-In cells were pre-treated ± AO (1 μM each) for 22 h prior to addition of TAK-243 for a further 2 h. Representative western blot and quantification of cleaved PINK1 (cPINK1) normalised to Actin shown relative to the TAK-243 (minus AO) condition. Error bars show standard deviation for three independent experiments. Note depolarisation induced cleavage (cl) of OPA1 in AO treated cells. **E.** hTERT-RPE1 YFP-Parkin cells were pre-treated ± AO (1 μM each) for 24 h prior to addition of TAK-243 (1 μM) for a further 4 h. Representative western blot of two independent experiments are shown. TOMM20 serves as a mitochondrial marker, which is lost after prolonged depolarisation in Parkin overexpressing cells. **F.** hTERT-RPE1 Flp-In PINK1 KO cells inducibly expressing PINK1-Flag were treated with doxycycline (Dox, 0.1 μg/ml) for 1 h ±AO (1 μM), ±TAK-243 (1 μM) as indicated and lysates subjected to immunoprecipitation with Flag-antibody. Representative western blot of three independent experiments and quantification showing cleaved pSer228-PINK1-Flag normalised to total cleaved PINK1-Flag. Error bars show standard deviation. FL: Full-length; cl: cleaved. **G.** Working model proposing two distinct pools of PINK1 on depolarised mitochondria: (i) a labile pool that is imported, processed and released as cleaved PINK1 from mitochondria, can be visualised (stabilised/trapped) by TAK-243 treatment; (ii) a separate pool of PINK1 remains intact and accumulates upon depolarisation. Both full-length and cleaved PINK1 are active in this model.

Phosphorylation at Ser228 provides a signature for PINK1 activation [18]. To determine if cPINK1 occurs in an activated form, we turned to an engineered RPE1 Flp-In cell line, in which endogenous PINK1 has been knocked out and replaced with an inducible PINK1-Flag transgene (Figure S1). This enabled us to immunoprecipitate PINK1-Flag and demonstrate its activity status using a pSer228-PINK1 directed antibody as a proxy reporter [18]. The level of cPINK1 accumulated upon TAK-243 treatment is independent of AO treatment, but its activation is strongly enhanced by such depolarisation (Figure 2F). Together these data suggest the co-existence of two distinct fates for PINK1, with one pool undergoing constitutive cleavage irrespective of membrane potential, whilst a second accumulates as a full length species only upon depolarisation. (Figure 2G).

### PINK1 stability measurements

Why does full length PINK1 accumulate under depolarising conditions, if the pathway leading to cPINK1 remains fully intact? Previous studies in Parkin over-expressing cells have suggested that this full length PINK1 accumulates in a stable form when treated with the protonophore CCCP [3]. Using a cycloheximide chase protocol, we have compared full length PINK1 half-life in untreated cells versus cells which have been treated for 4 hours with AO (Figure 3A). We take advantage of the high linearity and dynamic range of the LICOR Odyssey infrared scanner for quantitation, whilst also adjusting the protein loading levels between conditions to obtain comparable starting levels of PINK1. For untreated cells (basal state), the half-life of PINK1 is ∼30 minutes and to be at steady state, this degradation rate must be exactly balanced by the rate of protein synthesis (Figure 3B). Under AO depolarising conditions the pUb declines in step with PINK1 levels (Figure 3A). After 4 hours AO treatment PINK1 levels have increased 3.5 fold (Figure 3C), whilst mitochondrial membrane potential was substantively lost (Figure S2). However, only ∼25% of this PINK1 is in fact stable (Figure 3B and C), with the remainder showing a fast decay rate similar to untreated cells. This fraction of stable protein can only account for less than half of the increment we observe in PINK1 (Figure 3C). This accounting deficit cannot be ascribed to increased mRNA and is therefore most likely reflecting increased translation (Figure 3D and 3E).

**Figure 3.**
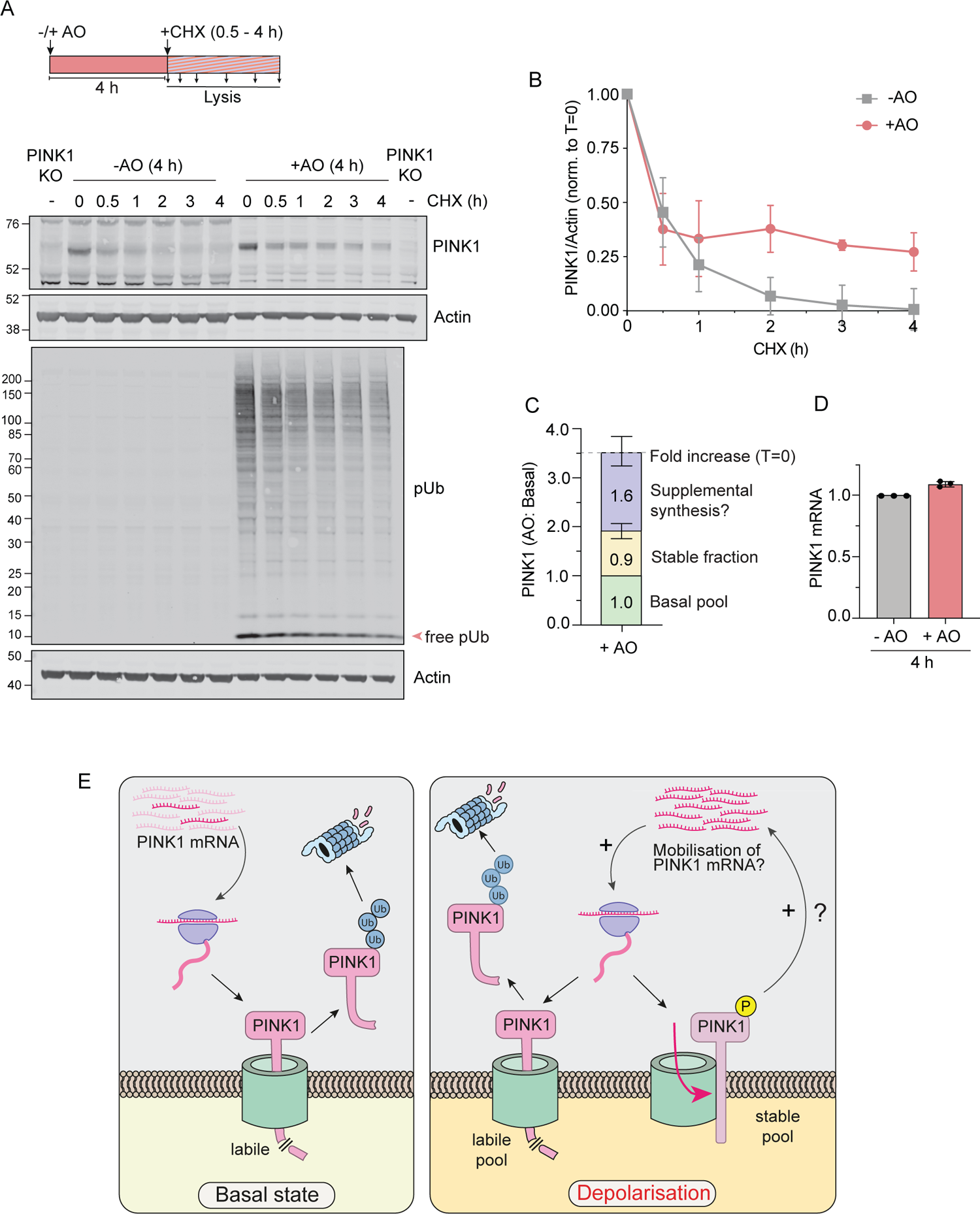
PINK1 stability measurements. **A.** hTERT-RPE1 Flp-In cells were treated for 4 h with AO (1 μM) followed by addition of cycloheximide (CHX; 100 μg/ml, ± AO) for 0.5 to 4 h. Experimental configuration and representative western blots are shown. Protein loading was adjusted to obtain a similar PINK1 signal at T=0 (3x protein loaded for the first 7 lanes in the PINK1 blot; 2x for the first 7 lanes of the pUb blot). An untreated PINK1 KO cell derived sample was loaded alongside for background subtraction. **B.** Quantification of the rate of PINK1 decay assessed as in A. Error bars show standard deviation, n=4 independent experiments. **C.** Graph showing the fold increase in PINK1 levels after 4 h of AO treatment (T=0h in A); the fraction of the accrued PINK1 that is stable (PINK1 (+AO) at T=3h/PINK1 (-AO at T=0h); and the proposed newly synthesised pool (Total fold increase - (basal pool + stabilised pool)). Error bars (fold increase and stable fraction) show standard deviation of four independent experiments. **D.** Quantitative RT–PCR reactions of PINK1 (normalised to Actin) were performed with cDNA derived from cells treated ± AO (1 μM) for 4 h. Error bars show standard deviation, n=3 independent experiments. **E.** Working model of PINK1 accumulation upon depolarisation. Under resting conditions (basal state), PINK1 is rapidly turned over through cleavage in mitochondria and subsequent ubiquitin-dependent degradation. This labile pool persists under depolarising conditions alongside the emergence of a stable pool of active PINK1, which is proposed to promote the mobilisation of PINK1 mRNA resulting in an increase in PINK1 protein synthesis.

### AO preconditioning of PINK1 activity

We conducted a series of experiments where we first subjected cells to a 4 hours treatment with AO and then washed out these drugs leading to re-establishment of the mitochondrial membrane potential (Figure S2). Firstly, TAK-243 was applied at the point of washout, and showed no influence on the dissipation rate of conjugated pUb (Figure 4A-C). Free pUb levels also decayed, suggesting that this rate is likely dominated by phosphatase activity. However, in TAK-243 treated samples we noticed a prominent second wave of free pUb generation, coincident with the emergence of cPINK1 (Figure 4B,D and E).

**Figure 4.**
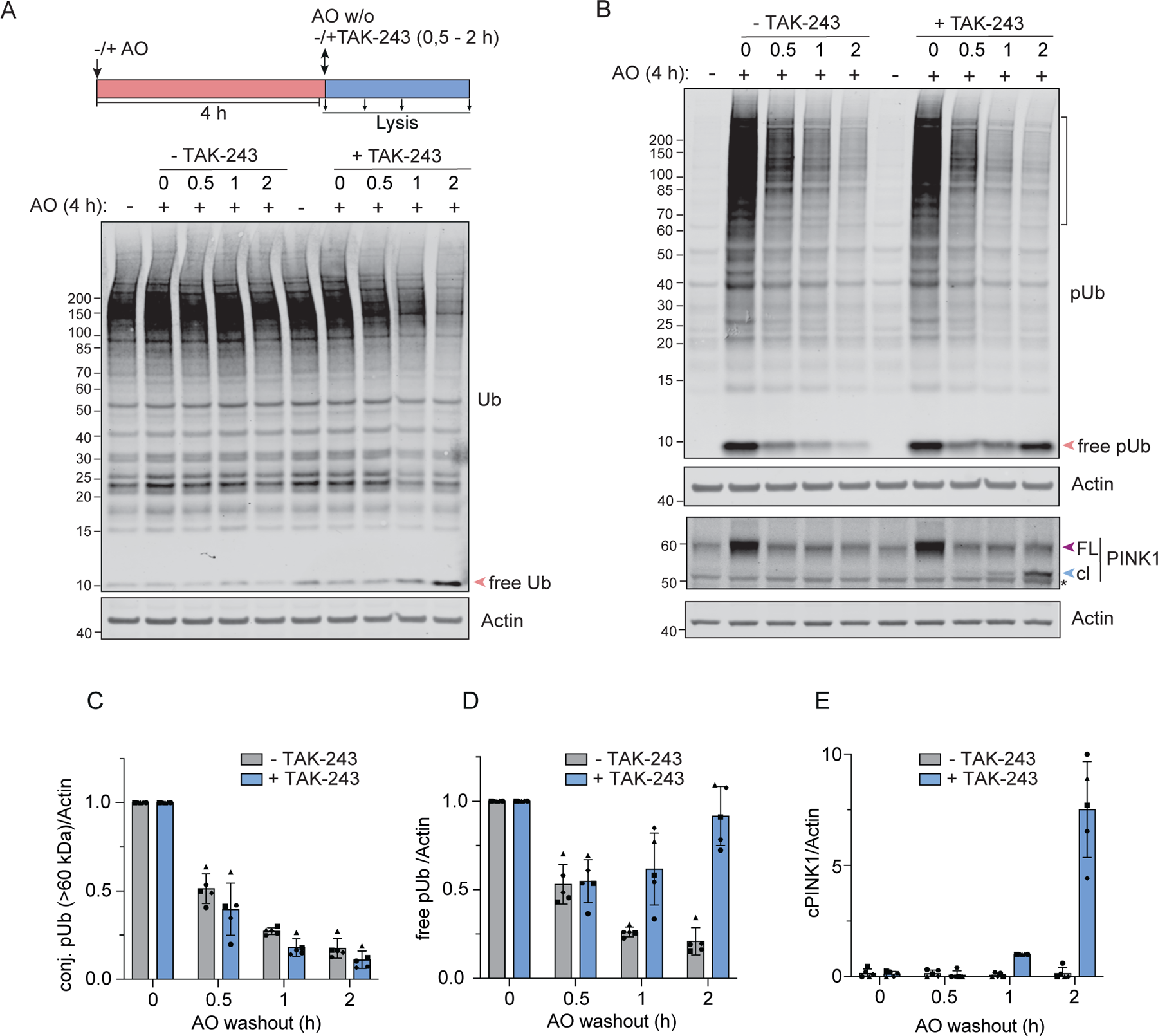
Correlation between cleaved PINK1 and free pUb generation upon E1 inhibition. **A.** Schematic diagram showing experimental overview and representative western blot for ubiquitin. hTERT-RPE1 Flp-In cells were treated ± AO (1 μM) for 4 h, followed by lysis (0 h) or replacement of media (washout) and a 0.5 to 2 h chase ± TAK-243 (1 μM). **B.** Representative Western blots for pUb and PINK1 from experiments depicted schematically in (A). The bracket indicates the range used for conjugated pUb quantification. * indicates a non-specific band. FL: Full-length PINK1; cl: cleaved PINK1. **C-E**. Quantification of data represented in (B) showing conjugated pUb (conj. pUb) decay (C), accumulation of free pUb (D) and cleaved PINK1 (cPINK1) (E). Error bars show standard deviation for 5 independent experiments.

We next modified our protocol to include an extended recovery time (16 hours) before the application of TAK-243. Remarkably, the AO-enabled activation of freshly generated PINK1 (cPINK1), is retained during this extended recovery period, even though the mitochondria fully recover their membrane potential within two hours (Figure 5A and Figure S2). We then turned to our doxycycline-inducible PINK1-Flag cell model to examine the activation status of PINK1 4 hours after TAK-243 administration following prior membrane depolarisation 16 hours beforehand (Figure 5B and C). As expected, we see evidence of cPINK1 activation, that is dependent on the prior mitochondrial depolarisation. A residual fraction of phosphorylated full length PINK1 is detectable, which may act as a seed for trans-phosphorylation of nascent PINK1 that is here being trapped as cPINK1 (Figure 5D). This is also consistent with the lingering shadow of a slightly upshifted endogenous PINK1 band, likely to be phosphorylated PINK1 that is maintained over the recovery period from AO treatment (Figure 5A, see lanes 3-7). These experiments reveal how such memory of a past event may be cached.

**Figure 5.**
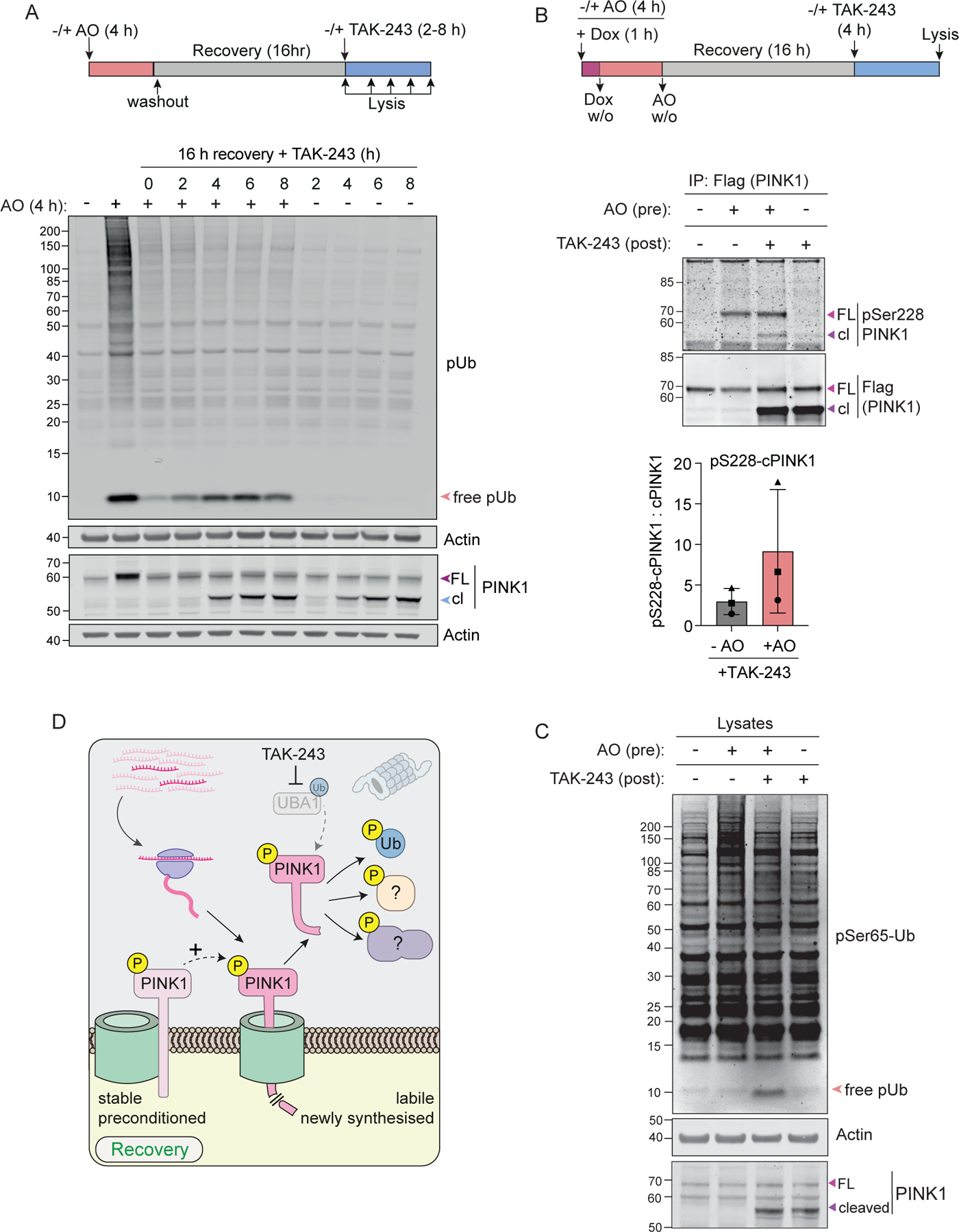
PINK1 exhibits sustained activity after mitochondria have recovered from depolarisation. **A.** Schematic diagram showing experimental overview and representative western blot for two independent experiments. hTERT-RPE1 Flp-In cells were treated for 4 h ± AO (1 μM) and either lysed immediately (lanes 1 and 2) or incubated in fresh media without AO (washout). After 16 h recovery, cells were either lysed (lane 3) or treated ± TAK-243 (1 μM) for an additional 2, 4, 6 or 8 h (lanes 4-11). **B and C**. hTERT-RPE1 Flp-In PINK1 KO cells inducibly expressing PINK1-FLAG were co-treated with doxycycline (Dox, 0.1 μg/ml) ± AO (1 μM) for 1 h, washed to remove Dox (w/o) and incubated ± AO in the absence of Dox for a further 3 h. AO was then washed out (w/o), and cells left to recover for 16 h before treatment ± TAK-243 (1 μM) for 4 h. Lysates were subjected to immunoprecipitation with Flag-antibody. Bound proteins (B) were analysed alongside the lysates (C) on SDS-PAGE. Western blots shown are representative of three independent experiments. Graph shows pSer228-cPINK1 normalised to total cPINK1 generated upon TAK-243 treatment after recovery from a prior depolarisation insult (AO). FL: Full-length PINK1; cl: cleaved PINK1. **D**. A small pool of PINK1 remains stable and active long after mitochondria have recovered from depolarisation, and can facilitate activation of nascent, labile PINK1, which can be visualised and stabilised by inhibiting ubiquitylation.

## Discussion

Acute application of the ubiquitin E1 conjugating enzyme inhibitor TAK-243 has allowed us to visualise the generation of cPINK1 [28]. The same product has previously been reported upon proteasome inhibition [3, 12, 32, 35]. However, TAK-243 treatment now offers an alternative method to generate this proteoform, with a divergent impact on the global distribution of ubiquitin. Both approaches provide an otherwise inaccessible glimpse of the constitutive mitochondrial cleavage rate of PINK1. Our finding that this rate is independent of polarisation conditions is unexpected given previous findings in HeLa cell models over-expressing Parkin [3]. The canonical model proposes that membrane potential driven insertion of the PINK1 pre-sequence into the the inner mitochondrial membrane (IMM) guided by TIM23 is necessary for exposure to the rhomboid protease PARL [36]. However, voltage independent turnover of PINK1 is not unprecedented. Experiments in HeLa cells lacking the small TOM complex subunit, TOM7, have revealed a corresponding cleavage under depolarising conditions, that is thought to be mediated by the depolarisation activated IMM protease OMA1 [17]. From this, it is suggested that TOM7 protects PINK1 from exposure to OMA1. Akabane *et al.* have also noted that protonophore-induced stabilisation of PINK1 is inhibited by coincident activation of Protein Kinase A, once again demonstrating efficient processing in the absence of a mitochondrial membrane potential [37].

How can we account for the observed accrual of full length PINK1 following depolarisation? We suggest that there are in fact two populations of newly synthesised PINK1; one which follows a voltage-insensitive pathway leading to degradation and a second voltage-sensitive pathway that leads to a stable pool of full length PINK1 (Figure 2G). We propose that all PINK1 interacts with a TOM complex for import, but that fates can differ according to the TOM channel status. The stable pool of PINK1 cannot account for the degree of accumulation we observe. Since we, and others, have also excluded an increase in PINK1 mRNA [3], we propose that PINK1 translation must increase under depolarising conditions. Previous global proteomic studies, have used the same logic to indicate the importance of differing translation efficiencies in determining protein abundance [38].

PINK1 mRNA is known to bind to mitochondrial membranes in neurons [26]. Similarly some respiratory chain component mRNAs can be recruited to mitochondria and are de-repressed in a PINK1 dependent manner [27]. The second effect of mitochondrial depolarisation is activation of PINK1 [4, 18, 39–41]. A simple speculative model posits that depolarisation leads to activation of the small fraction of pre-existing full length PINK1, which then acts to mobilise mitochondrial bound PINK1 mRNA for additional translation, which underlies the accrual of protein (Figure 3E).

Ischaemic preconditioning imparts tissue protection in multiple organs. This is a process whereby a short non-lethal ischaemia reperfusion protects against a subsequent severe ischaemia-reperfusion injury [42]. An intriguing study in the mouse kidney has identified mitophagy as a significant mediator of this response. Moreover following the first insult, PINK1 is upregulated and depletion of PINK1 leads to a loss of protection [43]. Here, we have discovered a conditioning phenomenon, directly linked to PINK1 activity. Treatment with AO conditions the response to subsequent application of TAK-243 many hours later. This leads to activation of nascent cleaved PINK1 and phosphorylation of free ubiquitin. Whilst this particular experimental condition has allowed us to discover this phenomenon, we think that the underlying mechanism may be relevant to more physiological settings that involve sequential insults to mitochondria. Our working model conceives that the medium for this “molecular memory” is provided by a small long-lived pool of activated PINK1, that primes the subsequent response (Figure 5D).

## Acknowledgements

We thank Jon Lane (University of Bristol) and Jonathon Pines (ICR, London) for providing the YFP-Parkin RPE1 and hTERT-RPE1 Flp-In TREX cell lines respectively, and Miratul Muqit (University of Dundee, UK) for the generous gift of anti-pSer228-PINK1 antibody. LP and IG have been funded by Wellcome Trust PhD studentships (102172/Z/13Z). ER-J has been funded by the Michael J Fox Foundation (MJFF-021278 18). MC is a recipient of a Royal Society Industry Fellowship INF\R2\212031.

## Materials and Methods

### Cell culture

hTERT-RPE1 Flp-In TREX (a gift from Jonathon Pines, London, UK) and hTERT-RPE1-YFP-PARKIN cells (a gift from Jon Lane, Bristol, UK) were cultured in Dulbecco’s modified Eagle’s medium DMEM-F12 supplemented with 10% FBS. hTERT-RPE1 Flp-In TREX cells expressing PINK1-3xFlag were maintained under Blasticidin (10 μg/ml) and G418 (400 μg/ml) selection and induced with Doxycycline (0.1 μg/ml). HCT116 cells were cultured in McCoy’s media supplemented with 10% FBS.

### Generation of PINK1 knockout (KO) cells

PINK1 KO cells were generated using CRISPR-Cas9 with a PINK1 targeting sgRNA (5’-CACCGTACCCAGAAAAGCAAGCCG-3’) cloned into pU6-(BbsI)-CBh-Cas9-T2A-mCherry (Addgene, #64324). This plasmid was transfected into hTERT-RPE1 Flp-In TREX cells followed 24 h later by fluorescence-activated cell sorting (FACS) to select for mCherry positive cells. Individual clones were isolated by single-cell dilution and validated by western blotting and genomic DNA sequencing. Clone 6 harboured a single base deletion (G44 in exon 1) in both alleles, resulting in a frameshift and downstream early stop codon at ORF position 106. This clone was used for further generation of PINK1-FLAG expressing cells.

### Generation of a hTERT-RPE1 TREX PINK1 KO (PINK1-Flag) cell line

C-terminally tagged PINK1-3xFLAG (PINK1-Flag) was cloned into pcDNA5 FRTTO NeoR by Gibson assembly and transfected into hTERT-RPE1 Flp-In TREX PINK1 KO cells alongside pOG44 (1:9). Two days after transfection, cells were transferred to Blasticidin (10 μg/ml) and G418 (400 μg/ml) containing media and grown under selection for two weeks. Individual clones were isolated by single-cell dilution and validated by immunofluorescence and western blot analysis. PINK1-FLAG expression was induced with 0.1 μg/ml doxycycline as indicated.

### Antibodies and reagents

Antibodies and other reagents are as follows: anti-PINK1 D8G3 (rabbit, 6946, 1:1000, Cell Signaling Technology), anti-pSer228-PINK1 (rabbit, generous gift from Miratul Muqit, MRC PPU, Dundee, UK, 1:170), anti-pSer65-Ub E2J6T (rabbit, 62802, 1:1000, Cell Signaling Technology), anti-UBA1 (rabbit, ab34711, 1:1000, Abcam), anti-UBC13 (rabbit, ab25885, 1:1000, Abcam), anti-Ubiquitin (mouse, VU101, 1:2000, LifeSensors), anti-p53 (mouse, sc-126, 1:1000, Santa Cruz), anti-OPA1 (rabbit, ab42364, 1:1000, Abcam), anti-Actin (mouse, 66009, 1:10000, Proteintech), anti-TOMM20 (rabbit, 11802-1-AP, 1:1000, Proteintech), anti-Flag (goat, NB600-344, 1:1000, Novus), anti-Flag (mouse, M2, F3165, 1:1000, Sigma-Aldrich), Cycloheximide (C7698, Sigma-Aldrich), Oligomycin A (75351, Sigma-Aldrich), and Antimycin A (A8674, Sigma-Aldrich), Doxycycline (D9891, Sigma-Aldrich), Epoxomicin (324800, Sigma-Aldrich), TAK-243 (MLN7243, S8341, Selleckchem), TMRE (ab113852, Abcam), Hoechst 33342 (62249, Thermo Fisher Scientific).

### Cell lysate preparation and Western Blotting

Cells were lysed on ice in NP-40 lysis buffer (0.5% Nonidet P 40 Substitute (74385, Sigma-Aldrich), 25 mM Tris–HCl, pH 7.5, 100 mM NaCl, and 50 mM NaF) supplemented with mammalian protease inhibitors (MPI) (Sigma-Aldrich) and PhosSTOP (Roche). Proteins were resolved using 4-12% NuPage gels using MOPS or MES (pUb, total Ub) running buffers and transferred to 0.45 μm or 0.2 μm (pUb, total Ub) nitrocellulose membranes. Membranes were blocked in either 5% milk, 5% BSA (pUb, PINK1, pS228-PINK1) or 0.1% fish skin gelatin (Ub) and 0.1% Tween20 in TBS prior to primary antibody incubation in the same blocking solution. Western blot visualisation was performed using IRdye 800CW and 680LT coupled secondary antibodies with an Odyssey infrared scanner (Li-COR, Biosciences). For quantitations, signal values were obtained from Image Studio software after subtracting the background signal.

### Subcellular fractionation

hTERT-RPE1 FLpIN TRex cells were washed with ice-cold PBS twice, harvested by scraping and centrifugation at 1000 g for 2 min at 4°C. Cell pellets were washed with HIM buffer (200 mM mannitol, 70 mM sucrose, 1 mM EGTA, and 10 mM HEPES–NaOH, pH 7.4) centrifuged again at 1000 g for 5 min and resuspended in HIM buffer supplemented with MPI (Sigma-Aldrich), PhosSTOP (Roche), and 50 mM 2-chloroacetamide (CAA; Sigma-Aldrich). Cells were mechanically disrupted by shearing through a syringe with a 27G needle. The resulting homogenate was cleared from nuclei and unbroken cells by centrifugation at 600 g for 10 min to obtain a post-nuclear supernatant (PNS). A crude mitochondrial fraction (Mito) was obtained by centrifugation at 7,000 g for 15 min and the post-mitochondrial supernatant was further separated by centrifugation at 100,000 g for 30 min into light membrane (LM) and cytosolic fractions. The mitochondrial and light membrane fractions were resuspended in HIM buffer supplemented with MPI, CAA and PhosSTOP. Equal amounts of protein were loaded for each fraction.

### Immunoprecipitation

hTERT-RPE1 TREX PINK1 KO (PINK1-Flag) cells were treated with 0.1 μg/ml doxycycline to induce PINK1 expression and treated as indicated prior to lysis in SDS-Lysis buffer (2% SDS, 1 mM EDTA, 50 mM NaF) at 110°C. Lysates (700-3500 μg) were diluted 4-fold in IP dilution buffer (2.5% Triton X-100, 12.5 mM Tris pH 7.5, 187.5 mM NaCl) supplemented with MPI and PhosSTOP. Lysates were cleared by centrifugation and incubated overnight with 100 μl Pierce™ Anti-DYKDDDDK Magnetic Agarose (A36797). The immunoprecipitates were washed three times with IP wash buffer (1 part SDS-lysis buffer, 4 parts IP dilution buffer), twice with TBS, and once with Millipore water. Bound proteins were eluted with 50 μl of 1.5 x non-reducing sample buffer (4.5% SDS, 93.75 mM Tris-HCl pH 6.8, 15% glycerol) at 95°C for 10 min, then mixed with reducing sample buffer and analysed by SDS-PAGE.

### RNA isolation and qRT-PCR

RNA was extracted using the Qiagen RNA extraction kit (74106). cDNA was then generated using Thermo Scientific RevertAid H Minus reverse transcription (Thermo Fisher Scientific; 11541515) supplemented with RNasin (Promega; N2611), PCR nucleotide mix (Promega; U144B) and oligo (dT) 15 Primer (Promega; C1101). Quantitative PCR were performed in triplicate using primers against PINK1 (5’-GGACGCTGTTCCTCGTTA-3’; 5’-ATCTGCGATCACCAGCCA-3’) and ACTB (Actin, 5’-CACCTTCTACAATGAGCTGCGTGTG-3’; 5’-ATAGCACAGCCTGGATAGCAACGTAC-3’), and iTaq Mastermix (BioRad; 172–5171) in a BioRad CFX Connect real-time system. Fold change was calculated using the 2−ΔΔCt method.

### Mitochondrial membrane potential measurements

Cells were seeded into μ-dishes (Ibidi) and stained with 50 nM TMRE and 0.5 μg/ml Hoechst 33342 for 20 min before imaging at 37°C and 5% CO2 using a Zeiss LSM900 microscope and a 40x LD C-apochromat 1.1 NA water immersion objective. Images were processed and analysed using FIJI v2.0 for quantitation and Adobe photoshop for illustration. Graph shows the corrected total cell fluorescence (CTCF = integrated integrated density – (area of selected cell × mean fluorescence of background readings); number of cells imaged: DMSO, 49; 4 h AO, 70; 2 h recovery, 53; 16 h recovery, 57).

## Supplementary Figure legends

**Figure S1:**
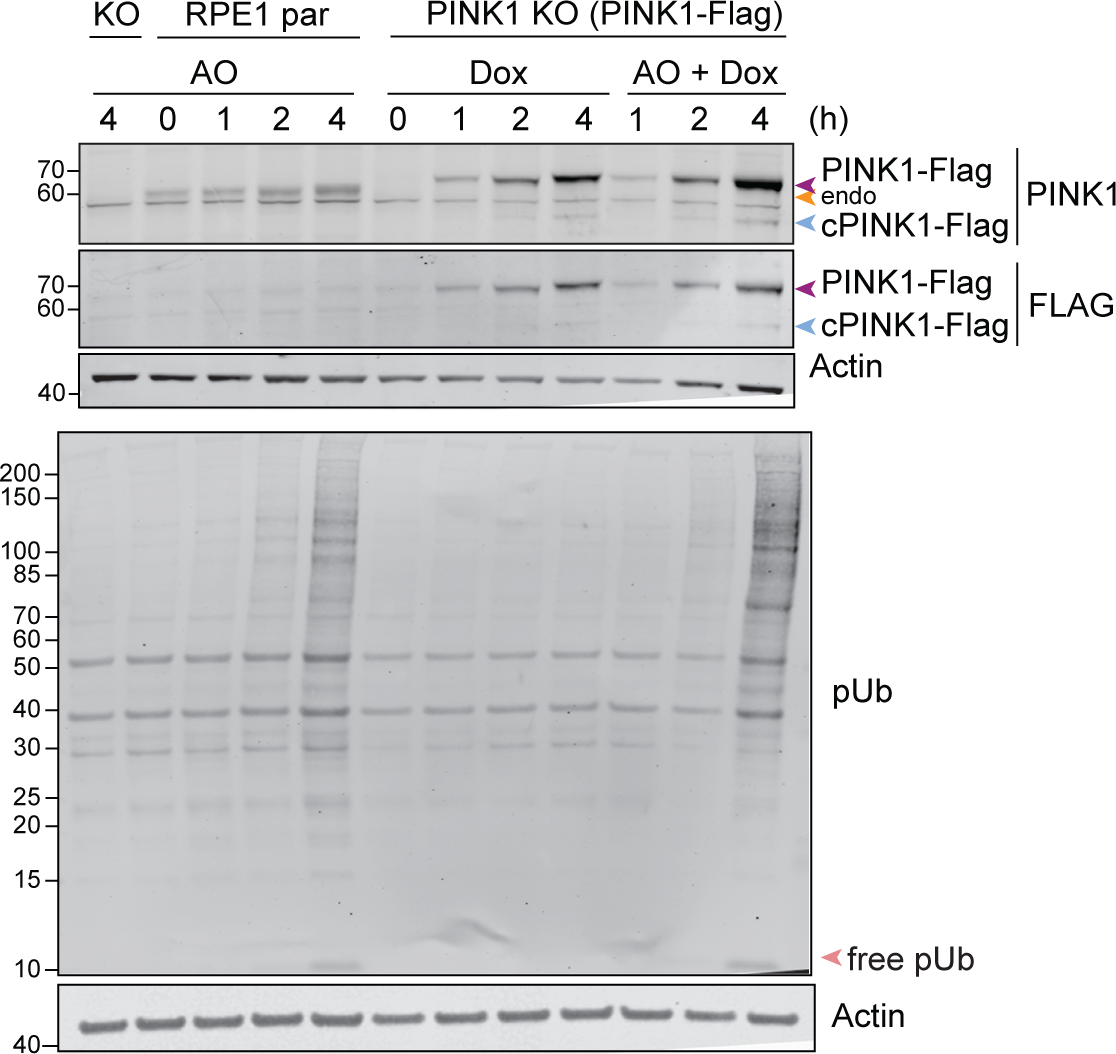
Characterisation of FL-PINK1-3xFlag expression in RPE1 PINK1 KO Flp-In FL-PINK1-3xFlag cells. **A.** hTERT-RPE1 Flp-In PINK1 KO cells (KO), parental hTERT-RPE1 Flp-In (RPE1 par), and hTERT-RPE1 PINK1-3xFLAG PINK1 KO cells were treated for the indicated time points ±Doxycycline (Dox, 0.1 μg/ml), ± Antimycin A and Oligomycin A (AO, 1 μM each) prior to lysis. Representative western blot of two independent experiments is shown, arrow head indicates endogenous (endo) PINK1.

**Figure S2:**
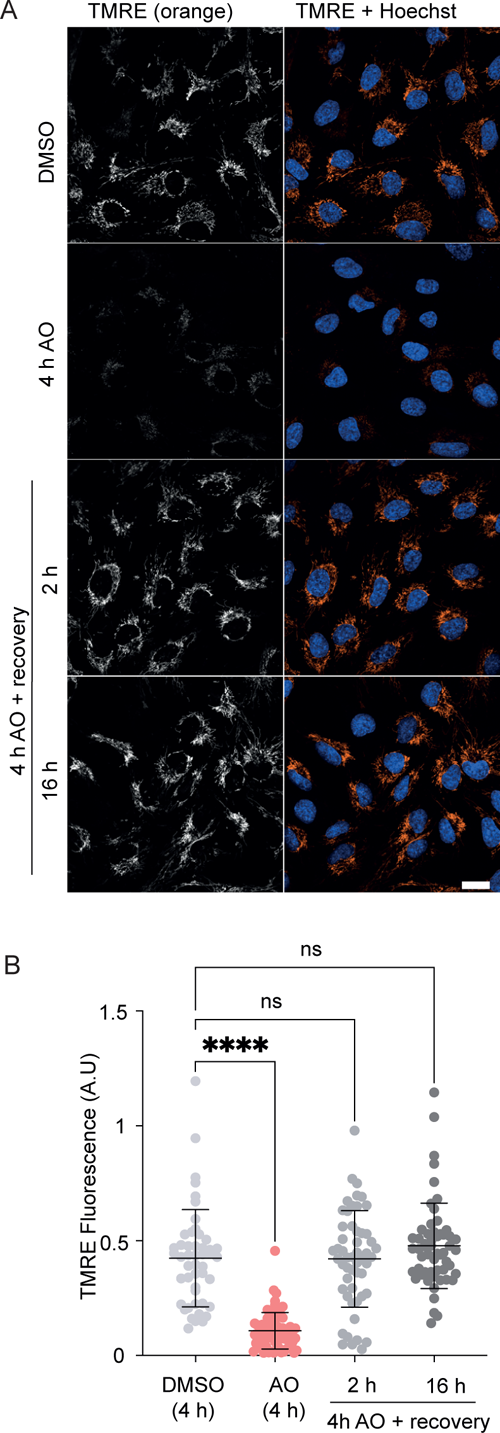
Analysis of mitochondrial membrane potential. **A.** hTERT-RPE1 Flp-In cells were treated ± AO (1 μM) for 4 h and either immediately stained with TMRE (50 nM) and Hoechst 33342 (0.5 μg/ml) or first allowed to recover for indicated timepoints (recovery) prior to staining. **B**. Quantification of TMRE fluorescence for each condition. Shown is the Mean Corrected Total Cell Fluorescence intensity for at least 49 cells per condition. Error bars indicate SD.

